# The vitamin B_3_ analogue nicotinamide riboside has only very minor effects on reducing muscle damage in *mdx* mice

**DOI:** 10.1101/2023.02.02.526793

**Authors:** Tiberiu Loredan Stan, Davy v.d. Vijver, Ingrid E.C. Verhaart, Annemieke Aartsma-Rus

## Abstract

**BACKGROUND:** Vitamin B3 analogue nicotinamide riboside (NR) has been suggested to have beneficial effects on muscle pathology in a mouse model for Duchenne muscular dystrophy (DMD). In muscle dystrophy, NR is thought to act acts by increasing levels of NAD+, to improve mitochondrial functioning and reduce muscle pathology.

**OBJECTIVE:** We here aimed to validate the effects of NR to improve muscle quality after eight weeks of treatment in two different mouse models for DMD: the commonly used mdx mouse on a C57BL/10 background (BL10*mdx*) and the more severely affected *mdx* mouse on a DBA/2J background (D2-*mdx*).

**METHODS:** To study in more detail whether NR treatment had an impact on muscle pathology, we assessed the expression levels of several markers for DMD pathology (fibrosis, regeneration and inflammation) in diaphragm.

**RESULTS:** Our data showed a trend for increase in NAD^+^-levels in blood; only in the D2-*mdx* NR-treated mice the NAD^+^-levels were slightly increased. These markers were elevated in *mdx* models compared to controls, but not affected by the NR treatment. Histological analysis of muscle tissues indicated a mild treatment effect in D2-*mdx* mice.

**CONCLUSIONS:** Based on our results, testing NR treatment in clinical trials in DMD patients is not warranted.

## INTRODUCTION

Duchenne muscular dystrophy (DMD) is an inherited, severe, progressive muscle wasting disorder with an incidence of around 1 in 5000 new-born boys [1]. In early childhood, patients show delayed development of motor function and signs of muscle weakness. Muscle function rapidly declines, leading to loss-of-ambulation around the age of ten, upon which respiratory and cardiac failure develops, leading to death, mostly in the third decade of life [2]. It is caused by genetic defects in the *DMD* gene, leading to the absence of a functional dystrophin protein, which plays an important role in stabilising muscle cells [3, 4]. Furthermore, its absence leads to disturbances in many other processes, like inflammation, fibrosis, regeneration and oxidative stress [5].

Life expectancy has greatly increased over the past years, mainly due to improved care and the use of corticosteroids [6], but treatment options stay limited. So far, only five drugs aiming to (partly) restore dystrophin protein expression are on the market, which have low efficiency and are applicable only to a subset of patients. Research on gene therapy is ongoing, but none has been approved yet [7]. Therefore, drugs targeting secondary defects are valuable. In addition, primary approaches will only restore a partially functional dystrophin protein and their efficacy relies, among others, on the quality of the muscle tissue, further keeping therapies targeting pathological pathways highly relevant [8].

Several of the secondary consequences of the absence of dystrophin are related to mitochondrial dysfunction [9, 10]. This is, among others, caused by increased production of reactive oxygen species (ROS; oxidative stress) and increased intracellular Ca^2+^ levels [11]. Some drugs targeting these defects have been tested in clinical trials, like idebenone [12, 13], metformin combined with L-citrulline or L-arginine [14] and epigallocatechin gallate (green tea extract) [15]. In dystrophic muscle fibres nicotinamide adenine dinucleotide (NAD^+^) levels are decreased, while the activity of the NAD^+^ consuming PARP protein is increased [16-18]. These decreased NAD^+^ levels cause a disbalance in the NAD^+^/NADH ratio, which is monitored by the nutrient sensor SIRT1. The disbalance results in a metabolic fasting response, thereby causing disturbances in mitochondrial function [19, 20]. Increasing NAD^+^ levels has a positive effect on mitochondrial function [21, 22], among others by activating SIRT1, which in turn leads to activation of downstream targets like FoxO1, PGC-1α and PPARα [23-25]. In *mdx* mice, a mouse model for DMD [26], improving mitochondrial biogenesis by increasing SIRT1 with SIRT1 activators or crossing *mdx* mice with mice overexpressing SIRT1 due to a genetic modification, reduced muscle pathology and atrophy [17, 27-29]. Also, overexpression of SIRT1 or PGC-1α had a preventive effect on muscle atrophy and ameliorated the pathology in *mdx* mice [30-32].

Nicotinamide riboside (NR; a vitamin B_3_ analogue) is an NAD^+^ precursor, and leads to increased NAD^+^ levels [33-36]. Vitamin B_3_ analogues have shown beneficial effects in mitochondrial myopathy [20, 37] and ageing [38, 39]. Zhang *et al*. showed that one of the mechanisms whereby NR counteracts ageing is by delaying senescence of stem cells. In DMD, a depletion of muscle stem cells (MuSC) is seen, reducing the regenerative capacity. Therefore, NR may also have beneficial effects in DMD by improving the regenerative capacity of the MuSC and counteracting the mitochondrial dysfunction. Indeed, it has been shown that NR treatment of *mdx* mice prevented MuSC senescence, thereby improving regeneration [39]. Subsequent studies showed improved muscle function and reduced heart pathology in several DMD animal models. In *mdx* mice and *dys-1;hlh-1mutant Caenorhabditis elegans* (roundworm) reductions in inflammation and fibrosis, and protection against muscle damage were seen. Furthermore NR treatment could also induce functional improvement in the more severely affected *mdx/utrn*^*-/-*^ mouse [18].

NR is an attractive compound for therapeutic purposes. It is a relatively cheap, widely available compound. Administration is easy and patient-friendly (capsules taken orally). In addition, there is already knowledge on the safety and pharmacokinetic profile of NR in humans. These studies indicate NR is well-tolerated [38, 40-42] and can lead to increases blood NAD^+^ levels [43-46]. Currently trials for several conditions (*e*.*g*. cardiovascular diseases, mitochondrial dysfunction, ageing-related changes, chronic kidney disease and metabolic syndrome) are ongoing [47].

Here, we further investigated the effect of NR on muscle quality and several aspects of the disease pathology in two different mouse models for DMD.

## MATERIALS AND METHODS

### Animals and nicotinamide riboside treatment

Experiments were approved by the Animal Experiment Committee (Dierexperimentencommissie) of the Leiden University Medical Center (permit #13211) and executed following EU-guidelines. Mice were bred in the Experimental Animal Facility of the Leiden University Medical Center and were housed in individually ventilated cages at 20.5°C with 12 h of light/dark cycles and had *ad libitum* access to standard RM3 chow (SDS; Essex, UK) and water. Care was taken to minimise the burden and distress for the animals. *Mdx* mice on a C57BL/10ScSn (C57BL/10ScSn-DMD^*mdx*^/J; BL10-*mdx* [26]) or DBA2/J (D2.B10-Dmd^*mdx*^/J; D2-*mdx* [48]) background and their wild type counterparts (C57BL/10ScSn or DBA/2J) were used. Group sizes were based on previously observed expression levels of fibrotic markers in the diaphragm. To show a minimal difference in gene expression of 10%, a group size of eight mice was required. One wild type DBA/2J mouse died during the experiment, but this was unrelated to experimental procedures.

Nicotinamide riboside chloride (Elysium Health, Inc.; NY, USA) was dissolved in drinking water at a concentration of 3.5 mg/mL (12 mM) as described previously [49-51] and provided *ad libitum* in light protected bottles. NR was dissolved weekly and water was refreshed every 2/3 days. Treatment was started at four weeks of age and lasted for eight weeks. At the end of treatment, mice were fasted in the morning, whereupon mice where anaesthetised by isoflurane and blood was extracted via the eye. Mice were killed by cervical dislocation and muscles were isolated.

### Measurement of NAD^+^ blood levels

Prior to sacrifice, blood was collected from fasted mice in Greiner Bio One MiniCollect™ Tubes containing 3.2% sodium citrate (Thermo Fisher Scientific; The Netherlands). After gently mixing, blood was transferred to cryovials containing 0.5 M Perchloric acid acs reagent 70% (Thermo Fisher Scientific) and stored at -80°C until further processing. NAD^+^ in whole blood was quantified by Keystone bioanalytical (North Wales, UK) using an LC/MS/MS method. Sample preparation involved acid precipitation to isolated NAD^+^ from the blood. Isotopic labelled NAD^+^ was used as an internal standard for the quantitation (Calibration range 0.5-50 μg/mL). Extracted sample was applied to a C18 HPLC column connected with a Shimadzu 20AD HPLC system (Shimadzu; Tokyo, Japan) for the separation and quantified with a Sciex AI5000 mass spectrometer (Sciex; Framingham, MA, USA).

### Protein analysis

Protein was isolated from muscle samples by homogenizing the muscles in 1.4 mm Zirconium Beads prefilled tubes (OPS Diagnostics; Lebanon, USA), containing 100 mM Tris-HCl (pH 6.8) – 20% (w/v) sodium dodecyl sulphate (SDS), using a MagNA Lyser (Roche Diagnostics; Woerden, The Netherlands). Concentrations were measured using a Pierce bicinchoninic acid (BCA) protein assay kit (Thermo Fisher Scientific) according to manufacturer’s instructions. Fifty μg of total protein, heated for 5 min at 95°C, in a 75 mM Tris-HCl (pH 6.8), 15% (w/v) SDS, 20% (v/v) glycerol, 5% (v/v) β-Mercaptoethanol, and 0.001% (w/v) Bromophenol blue buffer was loaded on 1.0 mm thick Criterion XT Tris acetate (poly-acrylamide) gels with a linear resolving gel gradient of 4-12% (Biorad; Lunteren, The Netherlands). Gels were run at 75 V (0.07 A) for 1 hour and at 150 V (0.12 A) for 2 hours on ice. Proteins were blotted onto ready-to-use Trans-Blot Turbo transfer pack using the Trans-Blot Turbo system (BioRad) at 2.5 A (25 V) for 10 min, upon which membranes were blocked for 1 hour in buffer containing 5% non-fat dried milk powder (Elk; Campina; Amersfoort, The Netherlands). Membranes were washed one time in TBST buffer containing 10 mMTris-HCl (pH 8.0), 0.15 M NaCl and 0.005% (v/v) Tween20 and incubated overnight at 4°C in primary antibody. As primary antibodies rabbit-anti-SIRT1 (1:500; #PA5-17074; Thermo Fisher Scientific), mouse-anti-FoxO1 (1:500; #MA5-17078; Thermo Fisher Scientific), rabbit-anti-acetyl-FoxO1 (1:500; #PA5-104560; Thermo Fisher Scientific) and mouse-anti-beta-actin (1:5000; AC-15; Novus biologicals; Abingdon, UK) were used. Membranes were washed three times in TBST buffer, incubated for 1 hour at 4°C with secondary antibody, goat-anti-rabbit IgG (1:4000; IRDye 680LT; LI-COR Biotechnology; Bad Homburg, Germany) and goat-anti-mouse IgG (1:4000; IRDye 800CW; LI-COR Biotechnology), upon which they were washed twice in TBST and once in TBS. The Odyssey system and software (LI-COR Biotechnology) were used for visualization. Loading samples order for Western blot analysis BL10 background is provided in Table S1 and blotting images in Figure S1. Loading samples order for Western blot analysis DBA background is provided in Table S2 and blotting images in Figure S2.

### RNA isolation and quantitative PCR

Total RNA was extracted from muscle samples in 1.4 mm Zirconium Beads prefilled tubes (OPS Diagnostics, Lebanon, USA), containing TRIsure isolation reagent (GCBiotech; Waddinxveen, The Netherlands), using a MagNA Lyser (Roche Diagnostics). RNA was cleaned up with the NucleoSpin RNA II kit (Macherey-Nagel; Düren, Germany) according to the manufacturer’s instructions. Thereupon cDNA was synthesised using 400 ng of total RNA as a template, for which random N6 primers (Thermo Fisher Scientific) and Tetro reverse transcriptase (GCBiotech) were used, following manufacturer’s instructions. For qPCR, SensiMix reagents (GCBiotech) and the LightCycler 480 (Roche Diagnostics) were used, using a program consisting of 45 cycles of 95°C (10 s), 60°C (30 s) and 72°C (20 s). Gene expression levels were analysed by the LinReg qPCR method. Since no reference genes that were consistently expressed in both the C57BL/10ScSn- and DBA2/J-strain could be found, for the C57BL/10ScSn mice *Ap3d1* and *GapdH*, and for the DBA2/J mice *Hbms* and *Pak1ip1* were used for normalisation. Gene expression levels were determined for *Cd68, Col1a1, Ctgf, Lgals3, Loxp, Myh3, Myog* and *Stat3*. Primer sequences are provided in Table S3.

### Sirius Red staining

Fibrosis was examined by Sirius Red staining of diaphragm muscle sections. Directly after sacrifice muscles were isolated, embedded in optimal cutting temperature (OCT) compound (Tissue-Tek; Sakura Finetek; Torrance, CA, USA) and snap-frozen in liquid nitrogen cooled isopentane. Eight μm thick sections were cut with a cryostat (Leica CM3050 S Research Cryostat; Amsterdam, The Netherlands). Interleaving sections were collected for RNA and protein analysis. Collagen levels were quantified with Sirius Red staining. Sections were fixed for 10 min in 4% paraformaldehyde and for 5 min 100% ethanol. Sections, were air-dried, rinsed in deionised water and stained in Sirius Red solution (Sirius Red 3FB/Direct Red 80; Sigma-Aldrich; St. Louis, MO, USA). They were washed twice for 5 min in acetic acid water (0.5% Glacial Acetic Acid in deionized water), rinsed in deionized water and dehydrated using an ethanol gradient (80%-90%-100%). Sections were mounted in Pertex mounting medium (VWR International B.V.; Amsterdam, The Netherlands) and imaged using a BZ-X700 fluorescent microscope (Keyence; Mechelen, Belgium) at 10 times magnification and processed and stitched using BZ-X Analyzer (Keyence). After background correction using Adobe Photoshop CC 2018 (Adobe Systems Corporation; San Jose, CA, USA), the ImageJ Software (NIH) was used. Sirius Red positive areas were normalized to the total tissue area. Sections were examined by two examiners independently and the average of their results was used for analysis.

### Statistical analysis

For statistical analysis Prism 8 (GraphPad Software Inc., CA, USA) was used. Values are presented as means ± standard deviation (SD). Histology, NAD^+^ levels, gene and protein expression were analysed with a two-way ANOVA between genotypes and age groups with one-way ANOVA followed by Tukey’s multiple comparison test to correct for multiple testing. In case of unequal variance a non-parametric Kruskal-Wallis test with Dunn’s post hoc correction was used. Statistical significance was set at *p*<0.05.

## RESULTS

### NAD^+^ levels in BL10-mdx mice and D2-mdx mice after nicotinamide riboside treatment

Firstly, we assessed the effect of NR in young *mdx* mice (four weeks of age) on a BL10 background after treatment for eight weeks. The treatment did not affect the bodyweight of the BL10-*mdx* mice (data not shown). At the end of treatment, NAD^+^ levels were assessed in the blood of treated and untreated mice and compared to their wild type counterpart (Fig. 1), with no significant changes for BL10-*mdx* mice (Fig. 1A). Although BL10-*mdx* mice have the same genetic mutation [52], they display a relatively mild phenotype compared to DMD boys [53]. This makes it more difficult to detect improvements and show therapeutic benefit of a test compound. Therefore, we also assessed the effect of NR in the more severely affected *mdx* mouse on a DBA/2J background (D2-*mdx*). At young age these mice exhibit severe symptoms of muscular dystrophy, like muscle atrophy and clear signs of fibrosis and inflammation. Furthermore, they have a lower body weight and functional performance is impaired [54-56]. Although the D2-*mdx* mice had a markedly lower bodyweight than their wild type counterparts, this parameter was not affected by eight weeks of NR treatment (start age four weeks; data not shown). The NAD^+^ blood levels were elevated in NR-treated D2-*mdx* mice, but this was only significant when compared to wild type DBA/2J mice (Fig. 1B).

**Figure 1.**
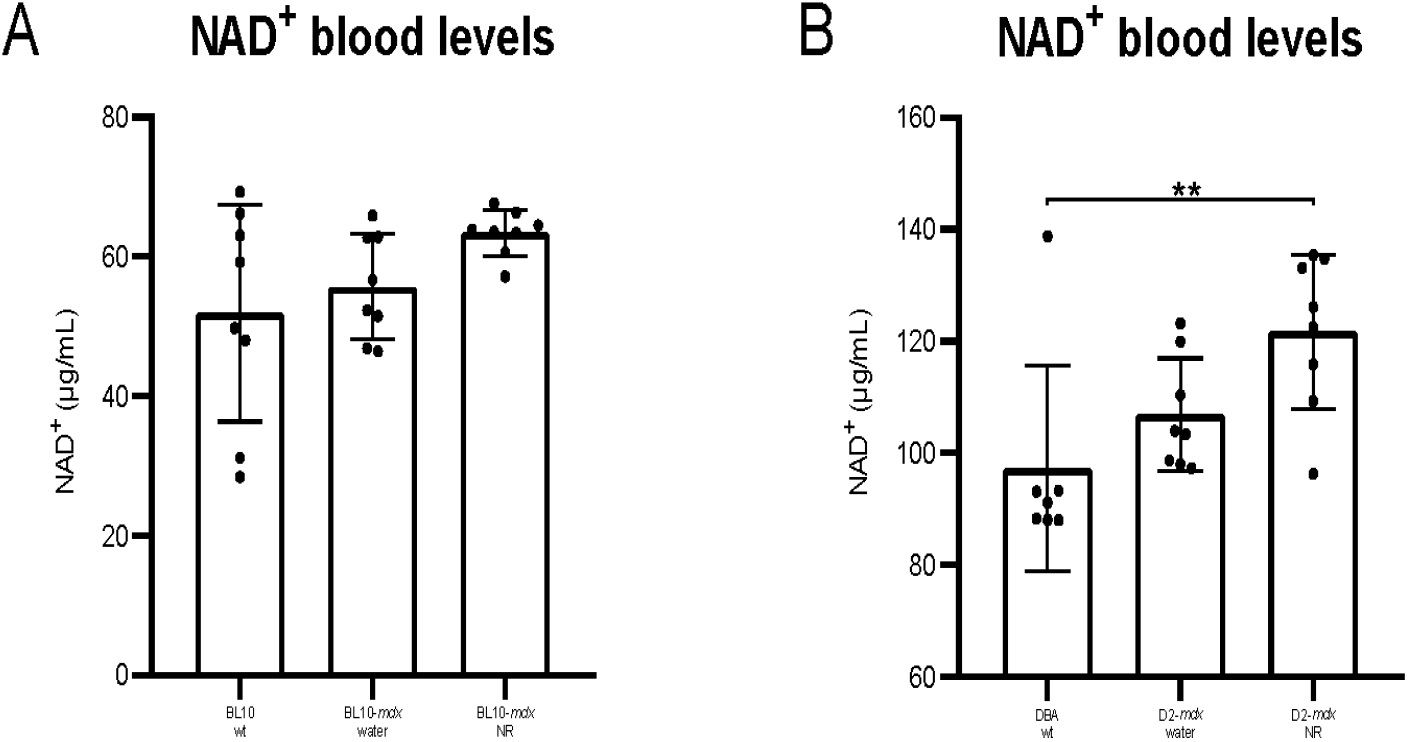
NAD+ blood levels in BL10-*mdx* (A) and D2-*mdx* (B) mice after 8 weeks of NR treatment. Due to differences in analysis, the levels of BL10-*mdx* and D2-*mdx* mice cannot be compared directly. Data represent mean ± SD (n=7-8 per group). One-way ANOVA followed by Tukey’s multiple comparison test.**p<0.01.

### Fibrosis

Since the diaphragm is the most severely affected muscle in mdx mice and thereby best resembles human pathology [57, 58], the diaphragm was used for all assessments. Fibrosis is one of the main characteristics of dystrophic muscle. The expression of two markers, lysyl oxidase (*Loxp*) and collagen, type Iα1 (*Col1a1*), was increased in BL10-*mdx* mice (Fig. 2A, B) although it could not be ameliorated by NR. Connective tissue growth factor (*Ctgf*) expression was not increased in BL10-*mdx* mice, nor was it affected by NR treatment (Fig. 2C). In D2-*mdx* mice disease pathology is characterised by fibrosis and inflammation, resulting in increased expression of several of gene markers. Similarly to BL10-*mdx* mice (Fig. 2A-C) an increase in the fibrotic markers *Loxp* and *Col1a1* was seen in D2-*mdx* mice and also here these were not decreased by NR treatment (Fig. 2D, E). In contrast to BL10-*mdx* mice (Fig. 2C) the *Ctgf* expression was increased in D2-*mdx* mice. Although a slight decrease in NR-treated D2-*mdx* mice could be observed, this was not statistically significant (Fig. 2D).

**Figure 2.**
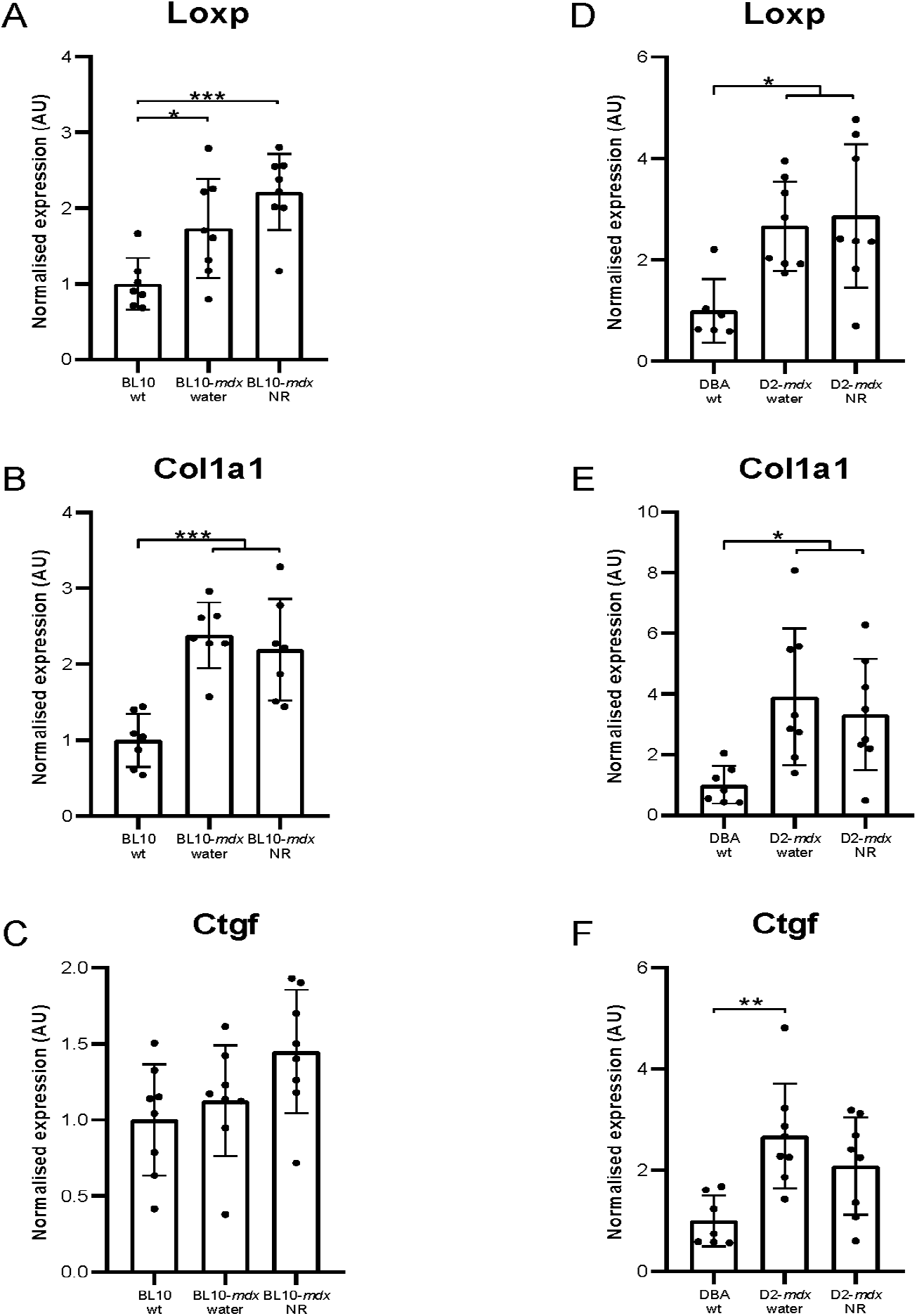
Fibrosis. Expression of fibrotic markers in BL10-*mdx* (A-C) and D2-*mdx* (D-F) mice. Due to differences in analysis, the levels of BL10-*mdx* and D2-*mdx* mice cannot be compared directly. Data represent mean ± SD (n=6-8). One-way ANOVA followed by Tukey’s multiple comparison test. *p<0.05, **p<0.01, ***p<0.001 compared to wild type mice.

### De- and regeneration

Muscular dystrophy causes massive de- and regeneration of muscle fibres as shown by increases in expression of the embryonic form of myosin heavy chain (*Myh3*) and myogenin (*MyoG*) in BL10-*mdx* mice (Fig 3A, B) and in D2-*mdx* mice (Fig. 3D, E). *Myh3* and *MyoG* are markers of early regeneration [59, 60] and were nearly undetectable in wild type mice (Fig. 3A-E). Signal transducer of transcription 3 (*Stat3*), a factor involved in satellite expansion and myogenic differentiation [61], was not increased in BL10-*mdx* mice (Fig. 3C), while this was seen in D2-*mdx* mice (Fig. 3F). None of these markers were influenced by NR treatment.

**Figure 3.**
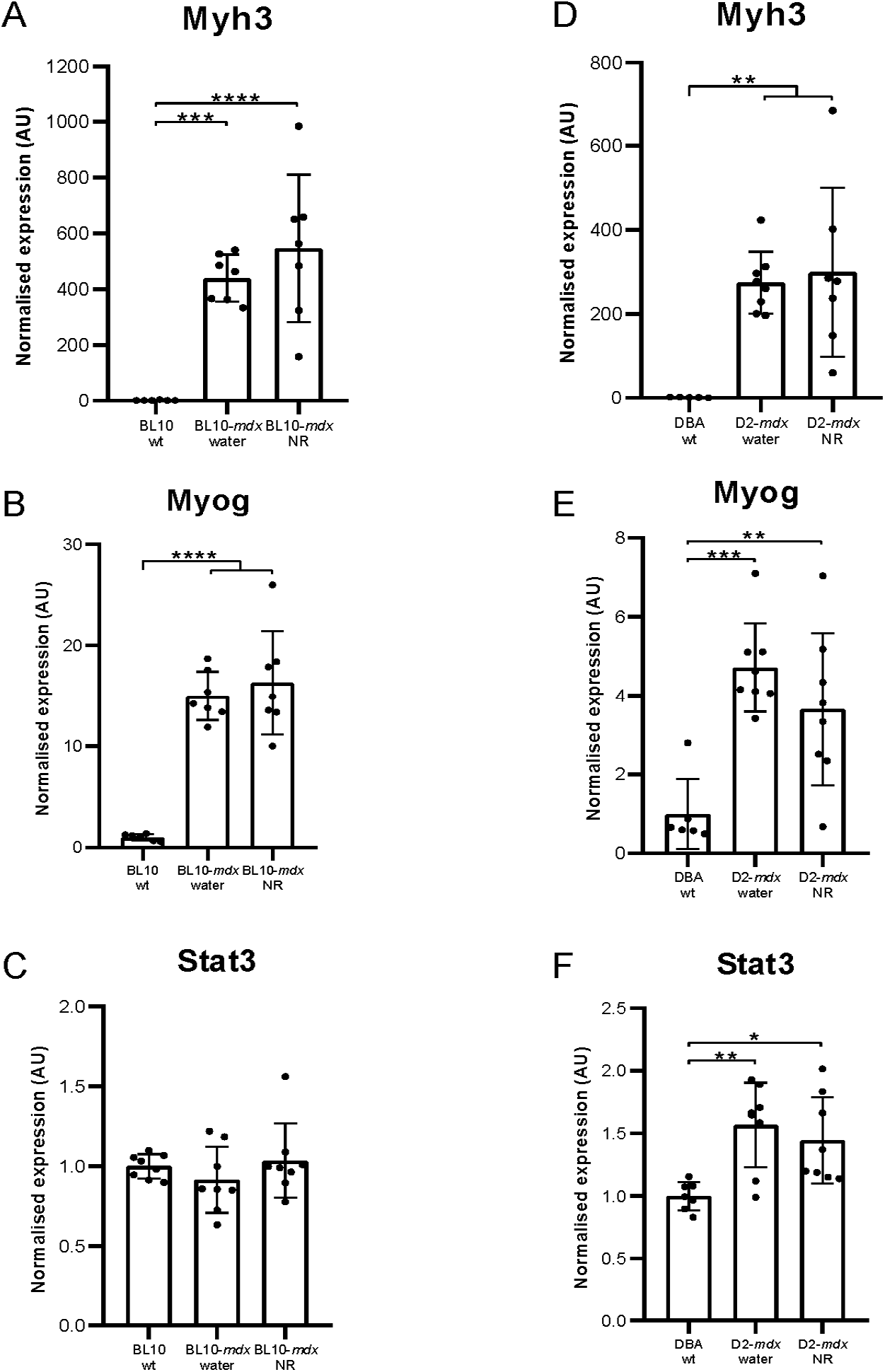
De- and regeneration. Expression of regenerative markers in BL10-*mdx* (A-C) and D2-*mdx* (D-F) mice. Due to differences in analysis, the levels of BL10-*mdx* and D2-*mdx* mice cannot be compared directly. Data represent mean ± SD (n=5-8). One-way ANOVA followed by Tukey’s multiple comparison test.*p<0.05, **p<0.01,. ***p<0.001 ****p<0.0001 compared to wild type mice.

### Inflammation

The breakdown of muscle tissue leads to inflammation. Indeed, large increases in both lectin, galactose binding, soluble 3 (*Lgals3*) and Cluster of differentiation factor 68 (*Cd68*) were seen in the BL10-*mdx* (Fig. 4A, B) and D2-*mdx* mice diaphragms (Fig. 4C, D), which were not decreased by NR treatment.

**Figure 4.**
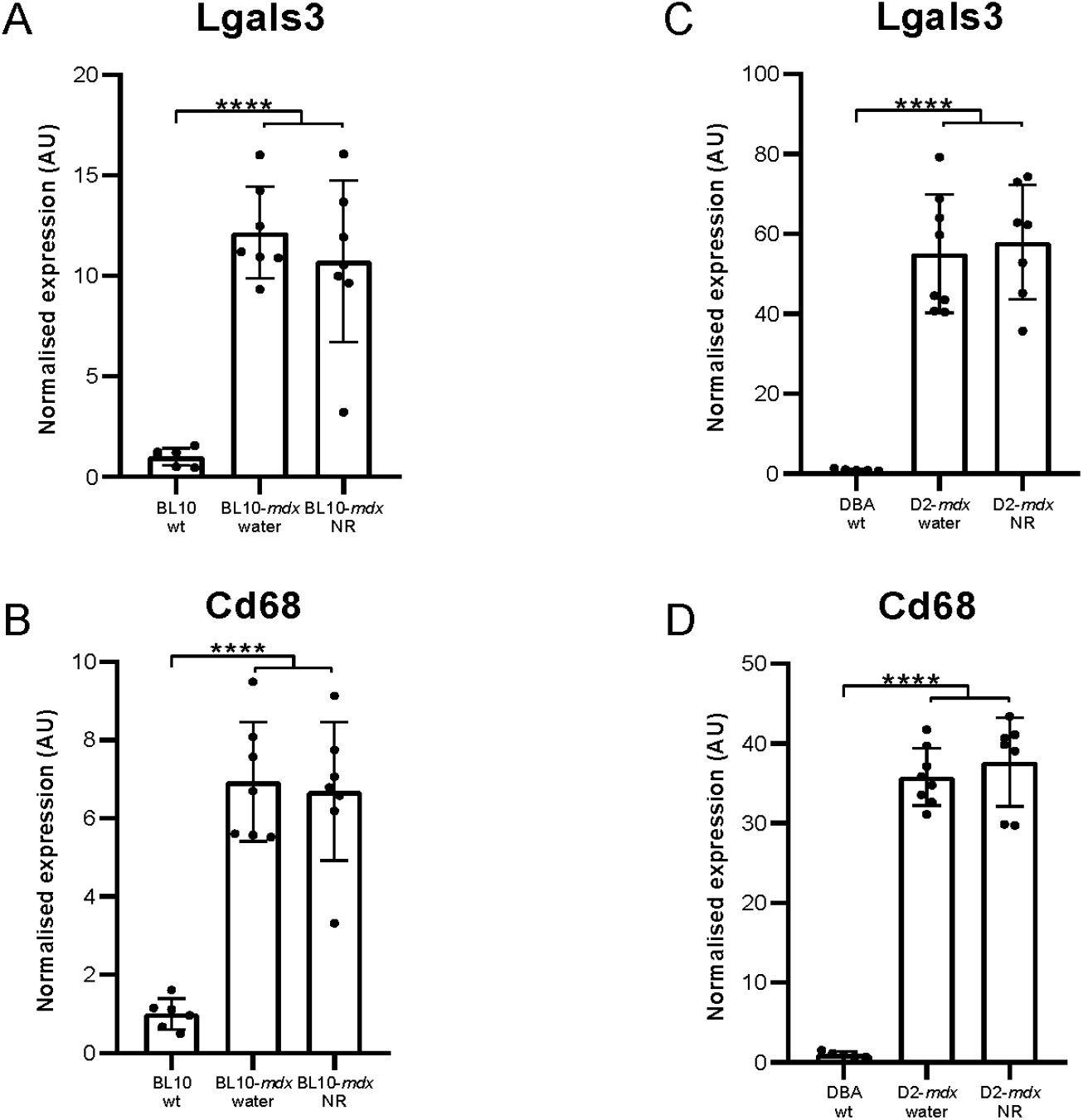
Inflammation. Expression of inflammatory markers in BL10-*mdx* (A-B) and D2-*mdx* (C-D) mice. Due to differences in analysis, the levels of BL10-*mdx* and D2-*mdx* mice cannot be compared directly. Data represent mean ± SD (n=5-8). One-way ANOVA followed by Tukey’s multiple comparison test. ****p<0.0001 compared to wild type mice.

### Muscle histology

In order to obtain a better characterisation of fibrosis throughout dystrophic degeneration we performed Sirius Red staining of the diaphragm sections in the considered mice (Fig. 5). Collagen content was significantly increased in water-treated BL10-*mdx* mice (Fig. 5A, B) and D2-*mdx* mice (Fig. 5C, D) when compared with their wild-type counterparts. NR treatment was not able to decrease the collagen accumulation neither for BL10-*mdx* nor for D2-*mdx* mice.

**Figure 5.**
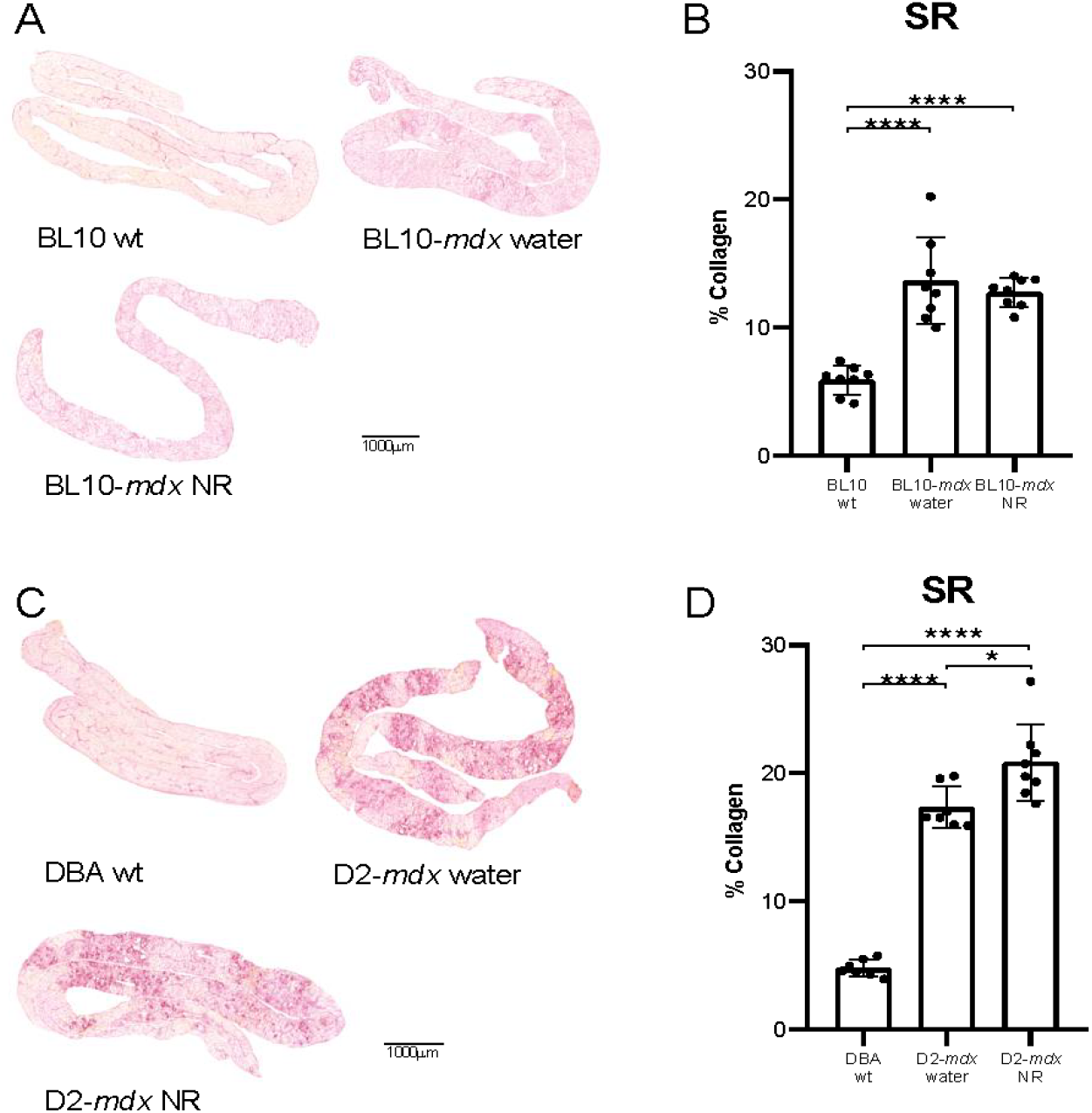
Muscle histology. Diaphragm of 12-week old BL10-*mdx* and D2-*mdx* mice after eight weeks of NR treatment. Representative images of Sirius Red stained diaphragms (A and C). Collagen tissue is stained red and muscle tissue yellow. Quantification of collagen content by Sirius Red staining (B and D) Data represent mean ± SD (n=7-8). One-way ANOVA followed by Tukey’s multiple comparison test. *p<0.05,****p<0.0001 compared to wild type mice.

### NAD+ downstream protein expression in diaphragm muscle

Increment of NAD^+^ leads to the activation of SIRT1 and its downstream targets, among others FoxO1 [23]. Therefore, we performed Western blot analysis to investigate the expression of downstream proteins (SIRT1, acetyl-FoxO1 and FoxO) in the diaphragm of BL10-*mdx* and D2-*mdx* mice (Fig. 6). For the for BL10-*mdx* mice the NR treatment did not significantly decrease the levels of SIRT1 (Fig. 6A), acetyl-FoxO1 (Fig. 6C) or FoxO (Fig. 6E). For the D2-*mdx*-NR treated mice the levels of SIRT1 (Fig. 6 B) and FoxO (Fig. 6F) did not change significantly when compared with D2-*mdx* water-treated mice. NR treatment however, decreased the level of acetyl-FoxO1 in D2-*mdx* mice (Fig. 6D).

**Figure 6.**
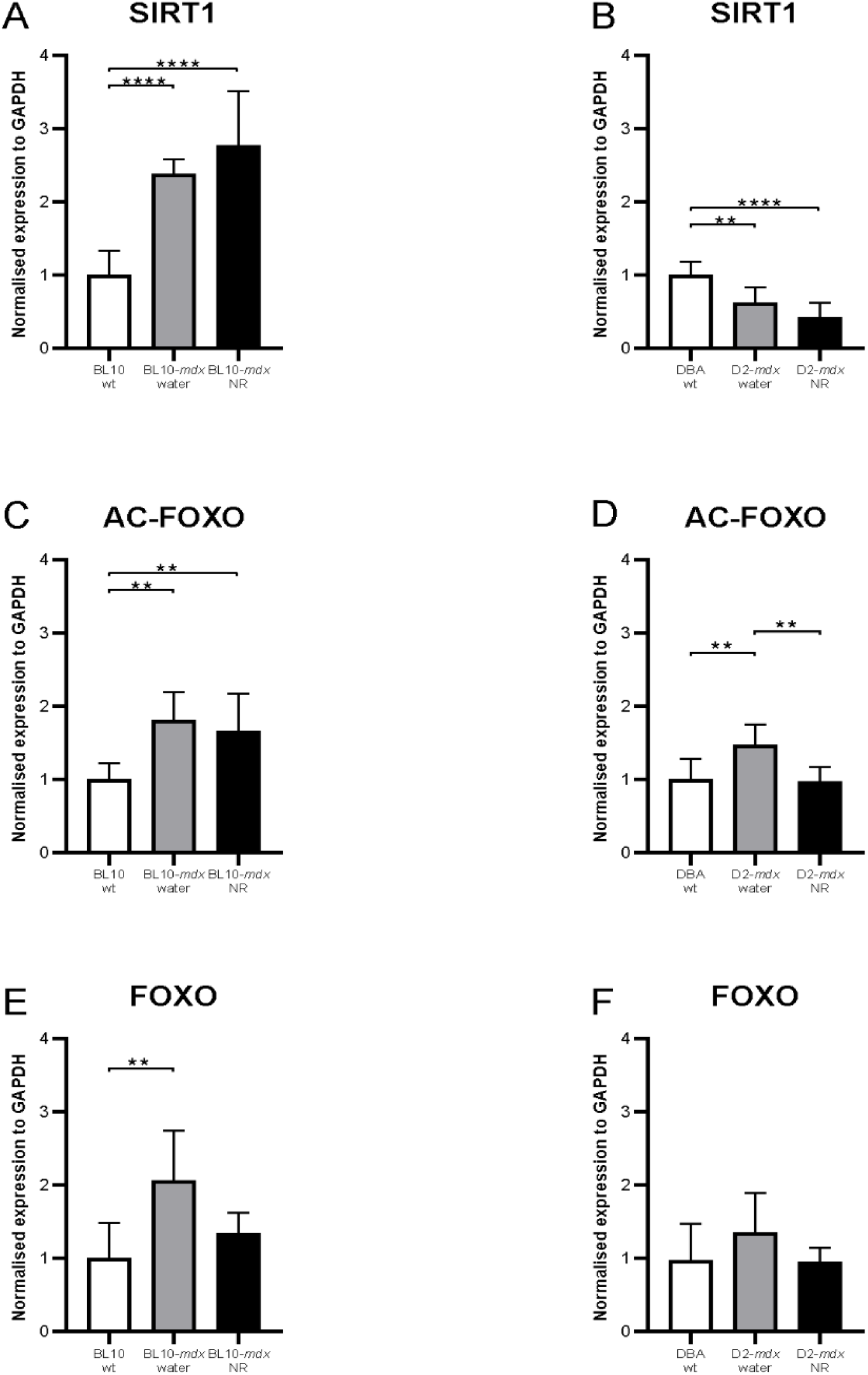
NAD+ downstream protein expression in diaphragm muscle. NAD+ downstream protein expression in diaphragm muscle of 12-week BL10-*mdx* (A, C, E) and D2-*mdx* (B, D, F) mice after eight weeks of NR treatment. Western blot analysis on expression of SIRT1 (A, B), acetyl-FoxO1 (C, D) and FoxO (E, F) normalized to GAPDH. Due to differences in analysis, the levels of BL10-*mdx* and D2-*mdx* mice cannot be compared directly mice. Data represent mean ± SD (n=7-8 per group). One-way ANOVA followed by Tukey’s multiple comparison test.**p<0.01, ****p<0.0001 compared to wild type.

## DISCUSSION

Although studies by the group of Auwerx *et al* showed promising results of NR in cell and animal models for DMD [18, 39] and NR treatment ameliorated disease pathology in models for several other conditions in which mitochondrial dysfunction plays a role [20, 37-39], we here could not demonstrate beneficial effects of NR in two mouse models for DMD. In both models clear signs of muscular dystrophy are observed, but there were no indications that NR could ameliorate disease pathology. One of the potential reasons for the discrepancies in the effect of NR between our study and those published may be the outcome parameters used. In particular, Ryu *et al*. reported that *mdx* mice show reduced NAD+ levels in skeletal hindlimb muscles, and that NR treatment improved mitochondrial and function (tibialis anterior), slightly reduced fibrosis in the diaphragm, reduced heart pathology and reversed pathology in *C. elegans*-DMD worm model. Zhang *et al*. reported that NAD+ repletion improves mitochondrial and stem cell function, modulates muscle stem cells senescence in *mdx* mice and improved the muscle function only in aged C57BL/6J mice. As such, NR might have an effect on some of aspects of muscular dystrophy and not on others.

Furthermore, although study set-up was largely comparable the route of administration of NR differed. In our study NR was dissolved in the drinking water (12 mM) instead of mixing it with the food (400 mg/kg/day). However, it is anticipated that this leads to comparable treatment amounts of NR per day. Furthermore, similar concentrations of NR in the drinking water as used in the presented study, has shown to be effective in other conditions [49, 51, 62]. In addition, a different type of NR was used. Here, we used NR chloride, while in the study of Ryu *et al*. NR triflate salt was used. We selected NR chloride because it is more clinically relevant, i.e. this is the type used by humans [63, 64]. It is possible that we do not see therapeutic effects as we did not achieve therapeutic increases in NAD^+^ levels. However, as stated before, we have used a concentration recommended and previously used by others [49, 51, 62] and increasing the NR dose is not an option due to tolerability issues. Except from a modest increase in the NR-treated D2-*mdx* mice, no effects on NAD^+^ levels itself were seen in our study and we also did not detect a decrease in NAD^+^ in both *mdx* strains compared to wild type mice. One explanation may be that these levels were measured in the blood and not in the muscle itself. However, if NAD^+^ levels in the muscle itself would have been affected by NR treatment, a change in expression in muscle of proteins in the downstream signalling pathway would be expected, which is not the case. Recently, several new studies [50, 65, 66] failed to show benefit of the NR treatment on DMD mouse models. Spaulding *et al*. looked at long term treatment of D2-*mdx* mice. After seven months of treatment no effects on muscle quality was seen nor were functional performance and respiratory functions improved [65, 66]. No additive effect of combination with several other nutraceuticals or pharmaceuticals previously shown to be beneficial for DMD were observed either. In this study, however, lower food concentrations of NR were used (55 mg/kg/day versus 400 mg/kg/day). Furthermore, the mice were treated at older age. Recent studies have shown that, although D2-*mdx* mice are severely affected at young age, pathology ameliorates with age [67]. This needs to be taken into consideration when planning experiments and may have prevented picking up a treatment effect as the difference between D2-*mdx* and wild types was much smaller than at a younger age. Furthermore, in a study by Frederick *et al*. in BL10-*mdx* mice no beneficial effect of 20-week treatment was seen. Although a decrease of NAD^+^ levels in muscle was seen in the mice after eccentric challenges, NR did not lead to an increment in these levels nor could it protect against eccentric injury. Inhibition of the NAD^+^ consuming enzymes ADP-ribosyl cyclases by the ecto-enzyme CD38 did result in increased NAD^+^ levels and functional benefits [50]. As such, the NR treatment does not robustly improve muscle pathology and function.

Most knowledge on the effect of NR on NAD^+^ metabolism is derived from rodent studies [18, 34, 39, 68]. Its effects in human skeletal muscle have less well been studied. Recent clinical trials in aged, overweight humans indicated that NR has no effect on the NAD^+^ levels itself, but does augment other metabolites of the skeletal NAD^+^ metabolome [38, 42]. One of the reasons for this may be the short half-life of NR in blood and that only limited amounts reach the muscles after oral ingestion [49, 69].

In conclusion, several studies show that increasing NAD^+^ levels is a promising therapeutic approach for DMD [17, 27-30, 50], but the vitamin B_3_ analogue NR is not the most efficient way to reach higher NAD^+^ levels and have beneficial effects in animal models for DMD.

## ACKNOWLEDGEMENTS

This study was supported by a grant from Duchenne UK. Nicotinamide riboside was kindly provided by Elysium Health, Inc. (NY, USA).

## CONTRIBUTION

IECV and AAR conceived and designed experiments. DvdV performed the experiments. IECV and AAR wrote the first draft of the manuscript. IECV, DvdV and TLS analyzed the data. TLS finalized the manuscript. All authors reviewed the manuscript and approved the final version.

## CONFLICT OF INTEREST STATEMENT

The authors declare no competing interest relevant to this study.

## SUPPLEMENTARY MATERIALS

Table S1. Loading sample order for Western blot analysis BL10 background

Table S2. Loading sample order for Western blot analysis DBA background

Table S3. Primer sequences used for gene expression analysis

Figure S1. Western blot images for analysis BL10 background. Different channels and loading samples order are depicted: RED:700 – GREEN:800, RAINBOW: 700 selection for SIRT and AC-FOXO and RAINBOW: 800 selection for FOXO and GAPDH.

Figure S2. Western blot images for analysis DBA background. Different channels and loading samples order are depicted: RED:700 – GREEN:800, RAINBOW: 700 selection for SIRT and AC-FOXO and RAINBOW: 800 selection for FOXO and GAPDH.

## Notes

### Competing Interest Statement

The authors have declared no competing interest.

### Summary of Updates

This version of the manuscript has been revised to include the figures in the main text.

