## Supplemental Table 1 for "The vitamin B_3_ analogue nicotinamide riboside has only very minor effects on reducing muscle damage in *mdx* mice"

**S1 Table. Loading sample order for Western blot analysis Bl10 background**

| **Blot 1** | **lower** |  |
| --- | --- | --- |
| Sample 1 | A1948496 | Wildtype C57BL/10 |
| Sample 2 | A1948500 | Wildtype C57BL/10 |
| Sample 3 | A1954203 | Wildtype C57BL/10 |
| Sample 4 | A1954204 | Wildtype C57BL/10 |
| Sample 5 | A1948464 | BL/10.*mdx* |
| Sample 6 | A1950334 | BL/10.*mdx* |
| Sample 7 | A1954164 | BL/10.*mdx* |
| Sample 8 | A1954657 | BL/10.*mdx* |
| Sample 9 | A1954163 | BL/10.*mdx* NR |
| Sample 10 | A1954658 | BL/10.*mdx* NR |
| Sample 11 | A1954660 | BL/10.*mdx* NR |
| Sample 12 | A1954731 | BL/10.*mdx* NR |
| **Blot 2** | **upper** |  |
| Sample 13 | A1954193 | Wildtype C57BL/10 |
| Sample 14 | A1954194 | Wildtype C57BL/10 |
| Sample 15 | A1956310 | Wildtype C57BL/10 |
| Sample 16 | A2001686 | Wildtype C57BL/10 |
| Sample 17 | A1954659 | BL/10.*mdx* |
| Sample 18 | A1954730 | BL/10.*mdx* |
| Sample 19 | A1954806 | BL/10.*mdx* |
| Sample 20 | A1954738 | BL/10.*mdx* |
| Sample 21 | A1954805 | BL/10.*mdx* NR |
| Sample 22 | A1954736 | BL/10.*mdx* NR |
| Sample 23 | A1954737 | BL/10.*mdx* NR |
| Sample 24 | A1955766 | BL/10.*mdx* NR |
| **upper gel sirt** |  |  |
| Image Name | Channel | Name |
| 0014053_02 | 700 | 1 |
| 0014053_02 | 700 | 3 |
| 0014053_02 | 700 | 4 |
| 0014053_02 | 700 | 5 |
| 0014053_02 | 700 | 6 |
| 0014053_02 | 700 | 7 |
| 0014053_02 | 700 | 8 |
| 0014053_02 | 700 | 9 |
| 0014053_02 | 700 | 10 |
| 0014053_02 | 700 | 11 |
| 0014053_02 | 700 | 12 |
| 0014053_02 | 700 | 13 |
| **lower gel sirt** |  |  |
| Image Name | Channel | Name |
| 0014053_02 | 700 | 14 |
| 0014053_02 | 700 | 15 |
| 0014053_02 | 700 | 16 |
| 0014053_02 | 700 | 17 |
| 0014053_02 | 700 | 18 |
| 0014053_02 | 700 | 19 |
| 0014053_02 | 700 | 20 |
| 0014053_02 | 700 | 21 |
| 0014053_02 | 700 | 22 |
| 0014053_02 | 700 | 23 |
| 0014053_02 | 700 | 24 |
| 0014053_02 | 700 | 25 |
| **upper gel Acetylated FOXO** | |  |
| Image Name | Channel | Name |
| 0014053_02 | 700 | 26 |
| 0014053_02 | 700 | 27 |
| 0014053_02 | 700 | 28 |
| 0014053_02 | 700 | 29 |
| 0014053_02 | 700 | 30 |
| 0014053_02 | 700 | 31 |
| 0014053_02 | 700 | 32 |
| 0014053_02 | 700 | 33 |
| 0014053_02 | 700 | 34 |
| 0014053_02 | 700 | 35 |
| 0014053_02 | 700 | 36 |
| 0014053_02 | 700 | 37 |
| **lower gel Acetylated FOXO** | |  |
| Image Name | Channel | Name |
| 0014053_02 | 700 | 38 |
| 0014053_02 | 700 | 39 |
| 0014053_02 | 700 | 40 |
| 0014053_02 | 700 | 41 |
| 0014053_02 | 700 | 42 |
| 0014053_02 | 700 | 43 |
| 0014053_02 | 700 | 44 |
| 0014053_02 | 700 | 45 |
| 0014053_02 | 700 | 46 |
| 0014053_02 | 700 | 47 |
| 0014053_02 | 700 | 48 |
| 0014053_02 | 700 | 49 |
| **upper gel GAPDH** | |  |
| Image Name | Channel | Name |
| 0014053_02 | 800 | 59 |
| 0014053_02 | 800 | 60 |
| 0014053_02 | 800 | 61 |
| 0014053_02 | 800 | 62 |
| 0014053_02 | 800 | 63 |
| 0014053_02 | 800 | 64 |
| 0014053_02 | 800 | 65 |
| 0014053_02 | 800 | 66 |
| 0014053_02 | 800 | 67 |
| 0014053_02 | 800 | 68 |
| 0014053_02 | 800 | 69 |
| 0014053_02 | 800 | 70 |
| **lower gel GAPDH** | |  |
| Image Name | Channel | Name |
| 0014053_02 | 800 | 71 |
| 0014053_02 | 800 | 72 |
| 0014053_02 | 800 | 73 |
| 0014053_02 | 800 | 74 |
| 0014053_02 | 800 | 75 |
| 0014053_02 | 800 | 76 |
| 0014053_02 | 800 | 77 |
| 0014053_02 | 800 | 78 |
| 0014053_02 | 800 | 79 |
| 0014053_02 | 800 | 80 |
| 0014053_02 | 800 | 81 |
| 0014053_02 | 800 | 82 |
| **upper gel FOXO** | |  |
| Image Name | Channel | Name |
| 0014053_02 | 800 | 83 |
| 0014053_02 | 800 | 84 |
| 0014053_02 | 800 | 85 |
| 0014053_02 | 800 | 86 |
| 0014053_02 | 800 | 87 |
| 0014053_02 | 800 | 88 |
| 0014053_02 | 800 | 89 |
| 0014053_02 | 800 | 90 |
| 0014053_02 | 800 | 91 |
| 0014053_02 | 800 | 92 |
| 0014053_02 | 800 | 93 |
| 0014053_02 | 800 | 94 |
| **lower gel FOXO** | |  |
| Image Name | Channel | Name |
| 0014053_02 | 800 | 95 |
| 0014053_02 | 800 | 96 |
| 0014053_02 | 800 | 97 |
| 0014053_02 | 800 | 98 |
| 0014053_02 | 800 | 99 |
| 0014053_02 | 800 | 100 |
| 0014053_02 | 800 | 101 |
| 0014053_02 | 800 | 102 |
| 0014053_02 | 800 | 103 |
| 0014053_02 | 800 | 104 |
| 0014053_02 | 800 | 105 |
| 0014053_02 | 800 | 106 |
| Checked all boxes for correct positioning | | |
| All boxes per protein have the same surface area | | |
