## Supplemental Table 2 for "The vitamin B_3_ analogue nicotinamide riboside has only very minor effects on reducing muscle damage in *mdx* mice"

**S2 Table. Loading sample order for Western blot analysis DBA background**

| **Blot 1** | lower |  |
| --- | --- | --- |
| Sample 1 | A2003216 | Wildtype DBA/2J |
| Sample 2 | A2003227 | Wildtype DBA/2J |
| Sample 3 | A2003228 | Wildtype DBA/2J |
| Sample 4 | A2003223 | Wildtype DBA/2J |
| Sample 5 | A1955802 | D2.*mdx* |
| Sample 6 | A1958648 | D2.*mdx* |
| Sample 7 | A1958650 | D2.*mdx* |
| Sample 8 | A2007265 | D2.*mdx* |
| Sample 9 | A1951247 | D2.*mdx* NR |
| Sample 10 | A1958649 | D2.*mdx* NR |
| Sample 11 | A1958651 | D2.*mdx* NR |
| Sample 12 | A2008588 | D2.*mdx* NR |
| **Blot 2** | upper |  |
| Sample 13 | A2007821 | Wildtype DBA/2J |
| Sample 14 | A2007822 | Wildtype DBA/2J |
| Sample 15 | A2007816 | Wildtype DBA/2J |
| Sample 16 | A2008589 | D2.*mdx* |
| Sample 17 | A2010329 | D2.*mdx* |
| Sample 18 | A2012978 | D2.*mdx* |
| Sample 19 | A2017167 | D2.*mdx* |
| Sample 20 | A2008590 | D2.*mdx* NR |
| Sample 21 | A2010328 | D2.*mdx* NR |
| Sample 22 | A2012162 | D2.*mdx* NR |
| Sample 23 | A2012977 | D2.*mdx* NR |
| **upper gel sirt** |  |  |
| Image Name | Channel | Name |
| 0014090_02 | 700 | 1 |
| 0014090_02 | 700 | 2 |
| 0014090_02 | 700 | 3 |
| 0014090_02 | 700 | 4 |
| 0014090_02 | 700 | 5 |
| 0014090_02 | 700 | 6 |
| 0014090_02 | 700 | 7 |
| 0014090_02 | 700 | 8 |
| 0014090_02 | 700 | 9 |
| 0014090_02 | 700 | 10 |
| 0014090_02 | 700 | 11 |
| **lower gel sirt** |  |  |
| Image Name | Channel | Name |
| 0014090_02 | 700 | 12 |
| 0014090_02 | 700 | 13 |
| 0014090_02 | 700 | 14 |
| 0014090_02 | 700 | 15 |
| 0014090_02 | 700 | 16 |
| 0014090_02 | 700 | 17 |
| 0014090_02 | 700 | 18 |
| 0014090_02 | 700 | 19 |
| 0014090_02 | 700 | 20 |
| 0014090_02 | 700 | 21 |
| 0014090_02 | 700 | 22 |
| 0014090_02 | 700 | 23 |
| **upper gel Acetylated FOXO** | |  |
| Image Name | Channel | Name |
| 0014090_02 | 700 | 24 |
| 0014090_02 | 700 | 25 |
| 0014090_02 | 700 | 26 |
| 0014090_02 | 700 | 27 |
| 0014090_02 | 700 | 28 |
| 0014090_02 | 700 | 29 |
| 0014090_02 | 700 | 30 |
| 0014090_02 | 700 | 31 |
| 0014090_02 | 700 | 32 |
| 0014090_02 | 700 | 33 |
| 0014090_02 | 700 | 34 |
| **lower gel Acetylated FOXO** | |  |
| Image Name | Channel | Name |
| 0014090_02 | 700 | 35 |
| 0014090_02 | 700 | 36 |
| 0014090_02 | 700 | 37 |
| 0014090_02 | 700 | 38 |
| 0014090_02 | 700 | 39 |
| 0014090_02 | 700 | 40 |
| 0014090_02 | 700 | 41 |
| 0014090_02 | 700 | 42 |
| 0014090_02 | 700 | 43 |
| 0014090_02 | 700 | 44 |
| 0014090_02 | 700 | 45 |
| 0014090_02 | 700 | 46 |
| **upper gel GAPDH** | |  |
| Image Name | Channel | Name |
| 0014090_02 | 800 | 47 |
| 0014090_02 | 800 | 48 |
| 0014090_02 | 800 | 49 |
| 0014090_02 | 800 | 50 |
| 0014090_02 | 800 | 51 |
| 0014090_02 | 800 | 52 |
| 0014090_02 | 800 | 53 |
| 0014090_02 | 800 | 54 |
| 0014090_02 | 800 | 55 |
| 0014090_02 | 800 | 56 |
| 0014090_02 | 800 | 57 |
| **lower gel GAPDH** | |  |
| Image Name | Channel | Name |
| 0014090_02 | 800 | 58 |
| 0014090_02 | 800 | 59 |
| 0014090_02 | 800 | 60 |
| 0014090_02 | 800 | 61 |
| 0014090_02 | 800 | 62 |
| 0014090_02 | 800 | 63 |
| 0014090_02 | 800 | 64 |
| 0014090_02 | 800 | 65 |
| 0014090_02 | 800 | 66 |
| 0014090_02 | 800 | 67 |
| 0014090_02 | 800 | 68 |
| 0014090_02 | 800 | 69 |
| **upper gel FOXO** | |  |
| Image Name | Channel | Name |
| 0014090_02 | 800 | 70 |
| 0014090_02 | 800 | 71 |
| 0014090_02 | 800 | 72 |
| 0014090_02 | 800 | 73 |
| 0014090_02 | 800 | 74 |
| 0014090_02 | 800 | 75 |
| 0014090_02 | 800 | 76 |
| 0014090_02 | 800 | 77 |
| 0014090_02 | 800 | 78 |
| 0014090_02 | 800 | 79 |
| 0014090_02 | 800 | 80 |
| **lower gel FOXO** | |  |
| Image Name | Channel | Name |
| 0014090_02 | 800 | 81 |
| 0014090_02 | 800 | 82 |
| 0014090_02 | 800 | 83 |
| 0014090_02 | 800 | 84 |
| 0014090_02 | 800 | 85 |
| 0014090_02 | 800 | 86 |
| 0014090_02 | 800 | 87 |
| 0014090_02 | 800 | 88 |
| 0014090_02 | 800 | 89 |
| 0014090_02 | 800 | 90 |
| 0014090_02 | 800 | 91 |
| 0014090_02 | 800 | 92 |
| Checked all boxes for correct positioning | | |
| All boxes per protein have the same surface area | | |
