## Supplemental Table 3 for "The vitamin B_3_ analogue nicotinamide riboside has only very minor effects on reducing muscle damage in *mdx* mice"

**S3 Table. Primer sequences used for gene expression analysis**

| **Gene** | **Full name** | **Primer** | **Sequence (5' - 3')** |
| --- | --- | --- | --- |
| ***Ap3d1*** | Adaptor-related protein complex 3 subunit delta-1 | forward | CTGAAGCAGGACAACATCGC |
|  |  | reverse | ATGATGTTGAAGGCAGCCCA |
| ***GapdH*** | Glyceraldehyde-3-phosphate dehydrogenase | forward | TCCCACTCTTCCACCTTCGA |
|  |  | reverse | CCACCACCCTGTTGCTGTAG |
| ***Hmbs*** | Hydroxymethylbilane synthase | forward | CCCGTAGCAGTGCATACAGT |
|  |  | reverse | ATGGTGGCCTGCATAGTCTC |
| ***Pak1ip1*** | P21-activated protein kinase-interacting protein 1 | forward | AAAGGAAGGTGGAGCATGGG |
|  |  | reverse | CGTCTTCTGCCCCACTGATT |
| ***Cd68*** | Cluster of differentiation factor 68 [1] | reverse | CTTCGGGCCATGTTTCTCT |
|  |  | reverse | AGAGGGGCTGGTAGGTTGAT |
| ***Col1a1*** | Collagen, type I, alpha 1 [2] | forward | ATGTTCAGCTTTGTGGACCT |
|  |  | reverse | CAGCTGACTTCAGGGATGT |
| ***Ctgf*** | Connective tissue growth factor [3] | forward | AGCTGGGAGAACTGTGTACG |
|  |  | reverse | GCCAAATGTGTCTTCCAGTC |
| ***Lgals3*** | Lectin, galactose binding, soluble 3 [4] | forward | CAACCATCGGATGAAGAACC |
|  |  | reverse | TTCCCACTCCTAAGGCACAC |
| ***Loxp*** | Lysyl oxidase [5] | forward | CAGAGGAGAGTGGCTGAAGG |
|  |  | reverse | CTGCCGCATAGGTGTCATAA |
| ***Myh3*** | Myosin, heavy polypeptide 3, skeletal muscle, embryonic [6] | forward | CGCAGAATCGCAAGTCAATA |
|  |  | reverse | CAGGAGGTCTTGCTCACTCC |
| ***Myog*** | Myogenin [7] | forward | CCCAACCCAGGAGATCATTT |
|  |  | reverse | GTCTGGGAAGGCAACAGACA |
| ***Stat3*** | Signal transducer of transcription 3 [8] | forward | GCTGCTGCATCTTCTGTCTG |
|  |  | reverse | TGAAGGTGGTGGAGAACCTC |
