## Supplementary figures and images for "The vitamin B_3_ analogue nicotinamide riboside has only very minor effects on reducing muscle damage in *mdx* mice"

### Supplemental Figure 1

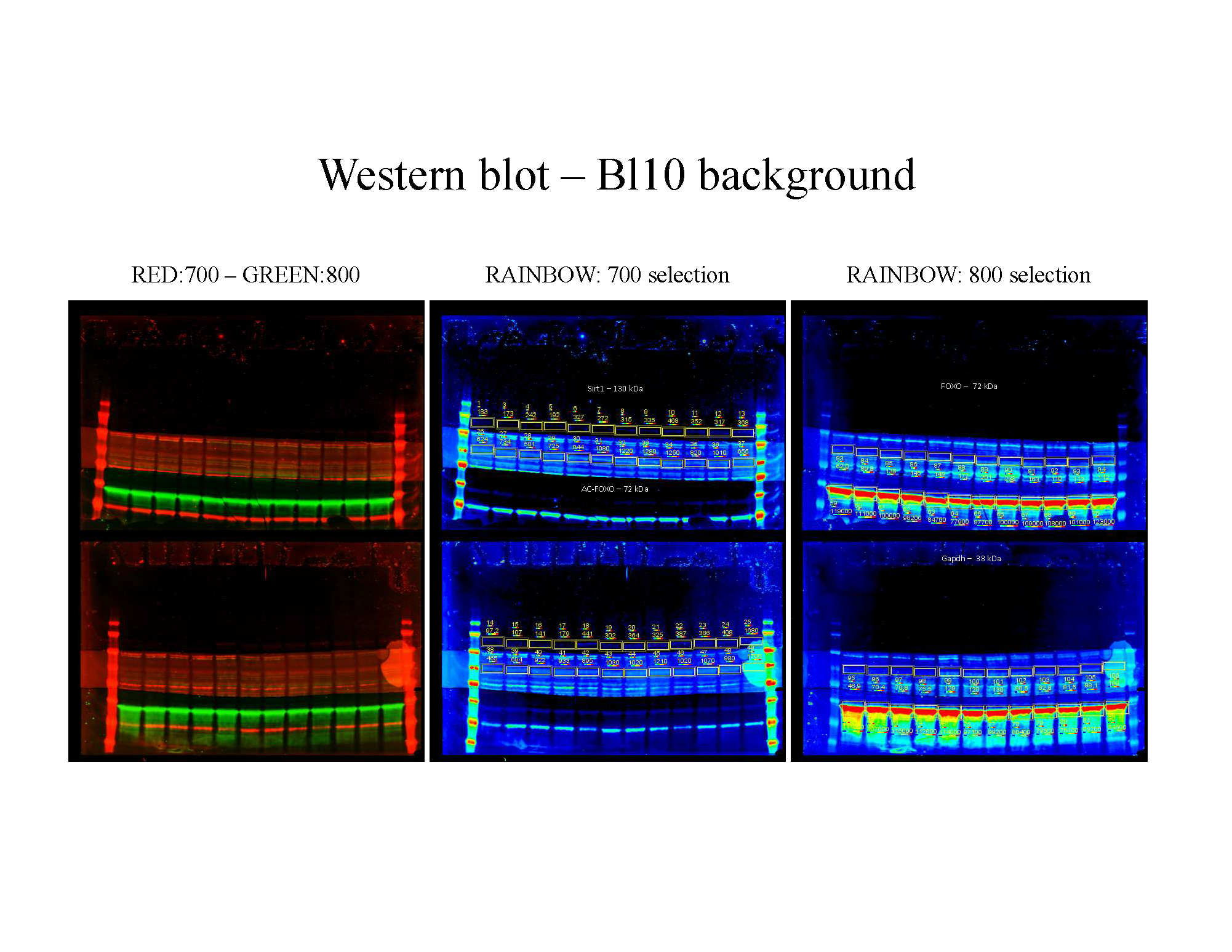

### Supplemental Figure 2

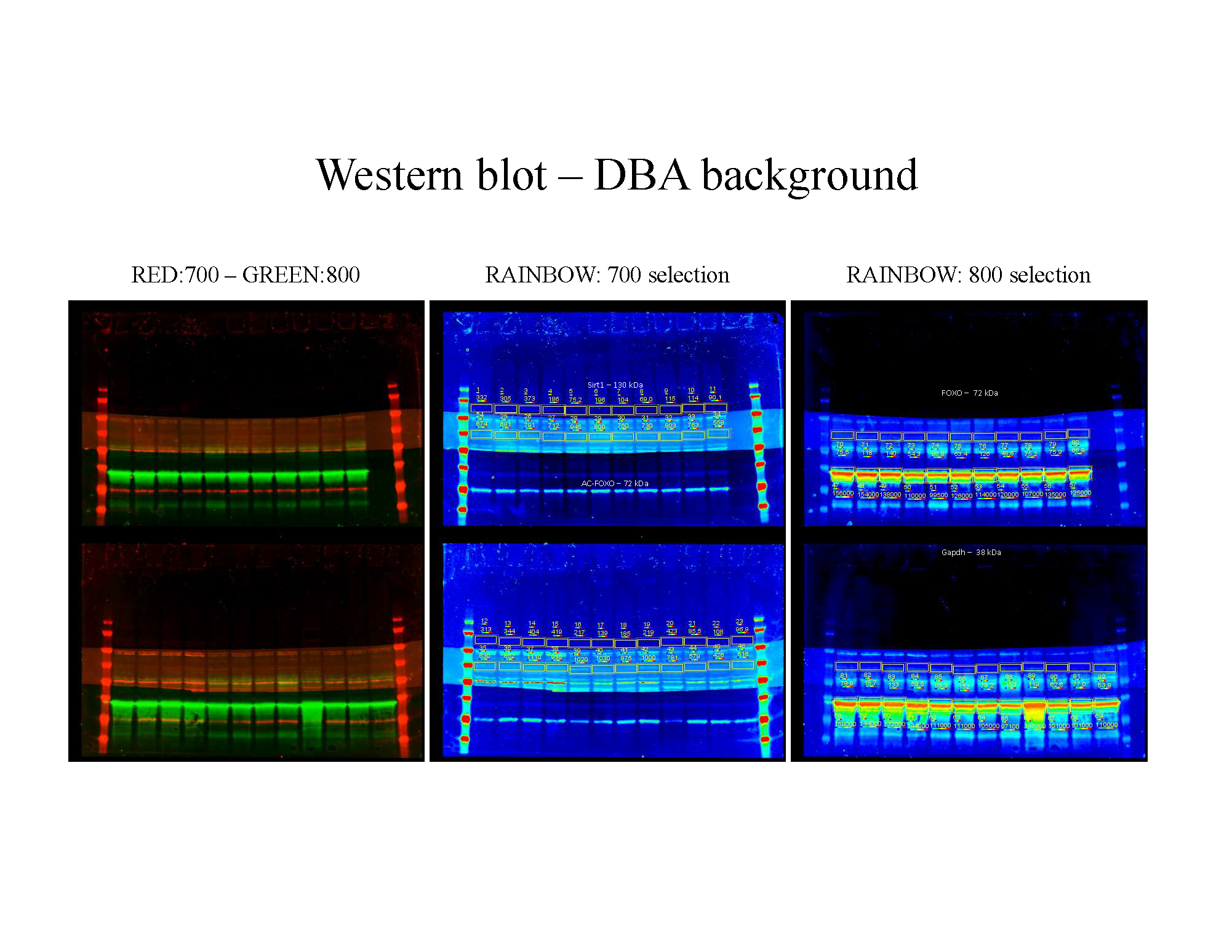
